# Bibliometric analysis of the trends of Zika related research from 2015 to 2017

**DOI:** 10.1101/287169

**Authors:** Yong-Dae Gwon, Magnus Evander

## Abstract

**Background:** Zika virus (ZIKV) is a mosquito-borne disease discovered in 1947, which did not cause public concern for the next 68 years. However, when ZIKV emerged in Brazil 2015 the attention increased rapidly. The announcement by Brazilian authorities, that ZIKV infection was associated with severe congenital disease e.g. microcephaly, surged public interest. Because of the accumulation of evidence that showed the magnitude of the ZIKV outbreak in the Americas, the World Health Organization declared a Public Health Emergency of International Concern February 1, 2016.

**Results:** During 2015-2017, we witnessed one of the most active and cooperated research responses against an emerging disease. To investigate the impact of ZIKV research during those years we decided to perform a bibliometric analysis of ZIKV research. The search for research articles on ZIKV was performed by bibliometric analysis from the scientific databases PubMed and Scopus. We found that the number of ZIKV related publications increased 38-41 times in 2016-2017 compared to 2015. During the three years there was a temporal shift in ZIKV research trends, from reports of ZIKV case studies and diagnostic methods, to development of ZIKV prevention and treatment. In addition, the number of countries involved in ZIKV research increased from 25 in 2015 to 111 in 2016 and 139 in the following year, showing that ZIKV research became global during three years.

**Conclusions:** The results from our study highlighted the importance of gathering public interest to global health issues, and how it can act as a powerful catalyzer to trigger the research field. However, despite the progress in ZIKV research, many questions remain to be addressed to accelerate the development of effective ZIKV countermeasures. Nevertheless, as long as we remember the importance of support and collaboration that we have experienced during the multidisciplinary effort against the current ZIKV outbreak, we will have an idea on how to handle the next inevitable and yet unknown infectious disease threat.

## Background

Zika virus (ZIKV) is a mosquito-borne virus in the family *Flaviviridae*, genus *Flavivirus*. Mainly mosquitoes from the *Aedes* genus are zoonotic vectors [1, 2]. After the first discovery 1947 in Uganda, Africa, ZIKV human infection has been found in Asia, the Pacific, and Americas [3, 4].

In 2007, a ZIKV outbreak was reported on Yap Island and nearby islands in Micronesia [5]. It later spread eastward in the Pacific Ocean and in 2015 it spread to the Americas with the initial identification in Brazil [6, 7]. After the 2015 introduction to the Americas, it was shown that ZIKV infection had a strong association to microcephaly in newborns, neurological disorders in adults (such as Guillain–Barré syndrome), as well as a potential for sexual transmission [8–10]. Due to the accumulation of evidence that showed the magnitude of the ZIKV outbreak in Americas, the World Health Organization (WHO) declared a Public Health Emergency of International Concern (PHEIC) on 1^st^ February 2016, and agreed on the urgent need to coordinate international efforts to investigate and understand this relationship better [11, 12].

According to the Organisation for Economic Co-operation and Development (OECD), bibliometrics is a statistical analysis of books, articles, or other publications to measure the “output” of individuals/research teams, institutions, and countries, to identify national and international networks, and to map the development of new (multi-disciplinary) fields of science and technology [13]. Scientific publication databases (e.g. Pubmed, Scopus, and Web of Science, etc) has been widely used for bibliometric analysis to assess the value of published scientific output in worldwide [14].

Therefore, we chose a bibliometric analysis method by using scientific publication databases to follow the trend of ZIKV research around the world. For analysis, we have collected publications for 2015-2017 and examined the trends of ZIKV research with various parameters.

## Methods

### Information sources and search strategy

To conduct the bibliometric analysis, we chose two different database sources, Pubmed and Scopus, for data extraction of publications related to ZIKV.

The search strategy was designed to find publication data about ZIKV from the scientific databases of published peer-reviewed studies. The search strings for the two different databases were selected from what has previously been used in a systematic review article from WHO [15]. The Pubmed search string was: zika [Title/Abstract] OR ZIKV [Title/Abstract] OR zika virus [MeSH Terms] OR zika virus infection [MeSH Terms]. The Scopus search string was: ZIKA [Title/Abstract/Keyword] OR ZIKV [Title/Abstract/Keyword] OR “zika virus” [Keyword] OR “zika virus infection” [Keyword].

We conducted our search for articles published from January 1^st^ 2015 to December 31^st^ 2017. Data for further analysis were selected and extracted, and then related to different parameters. All data were extracted April 2, 2018.

### Generation of the bibliometric summary report

Profiles Research Networking Software (Profiles RNS) is an NIH-funded open source tool to speed-up the process of finding researchers with specific areas of expertise for collaboration and professional networking [16]. The bibliometric summary report tool from Profiles RNS was used to calculate common metrics, including citation counts and h-index, from given a list of Pubmed IDs derived from Pubmed, a database of the U.S. National Library of Medicine by using Entrez Direct (EDirect) utility from a UNIX terminal window [17]. The generated report was analyzed for data sorting and represented as one of the parameters in this study.

### Data Extraction

To extract the ZIKV related publications for 2015-2017, we used Scopus with the search string mentioned above. The results were extracted in an Excel CSV document format. Next, we sorted out the result by analytic parameters (e.g. publication type, journal type, country, citation number, and core author), and the data used in this study is provided in the additional excel file (Additional file 1).

### Generation of geographic information system (GIS) maps

User-friendly GIS-maps [18], were generated to display the distribution of the publications. To generate and illustrate the GIS maps, we selected the R programming language with the “rworldmap” plug-in which is a package for visualizing global-scale data, concentrating on data referenced by country codes or gridded at half-degree resolution [19].

### Data visualization

All data visualization were performed by using GraphPad 7.0 (CA, USA). Adobe Illustrator CC 2017 (CA, USA) was used for editing visualized data.

## Results

### The ZIKV research field was fuelled by surged public interest

Until the signal flares alerted us about the new emergence of ZIKV in Americas, ZIKV was far from the public interest and poorly studied since 1947 [20]. Unexpectedly, the outbreak in the Americas was associated not only with mild infection, but also with severe congenital disease, particularly microcephaly in newborn infants. [21]

In addition, photographs that focused on a mother holding her newborn with microcephaly caused by ZIKV infection during pregnancy, caused sympathy and concerned authorities [22]. Once the public interest was raised, it led to a global awareness and actions for aiming to prevent the new ZIKV outbreak. Particularly in science, many research fields were fuelled to investigate and better understand all aspects of ZIKV.

We, as witnesses of one of the most rapid and organized research responses against an emerging disease, chose to compare the number of scientific publications to highlight the increase of research in the last three years. In 2015, there was an average number of 51 ZIKV related publication and the number then increased rapidly in 2016-2017 (38.3 and 40.6 times, respectively) (Figure 1).

**Figure 1.**
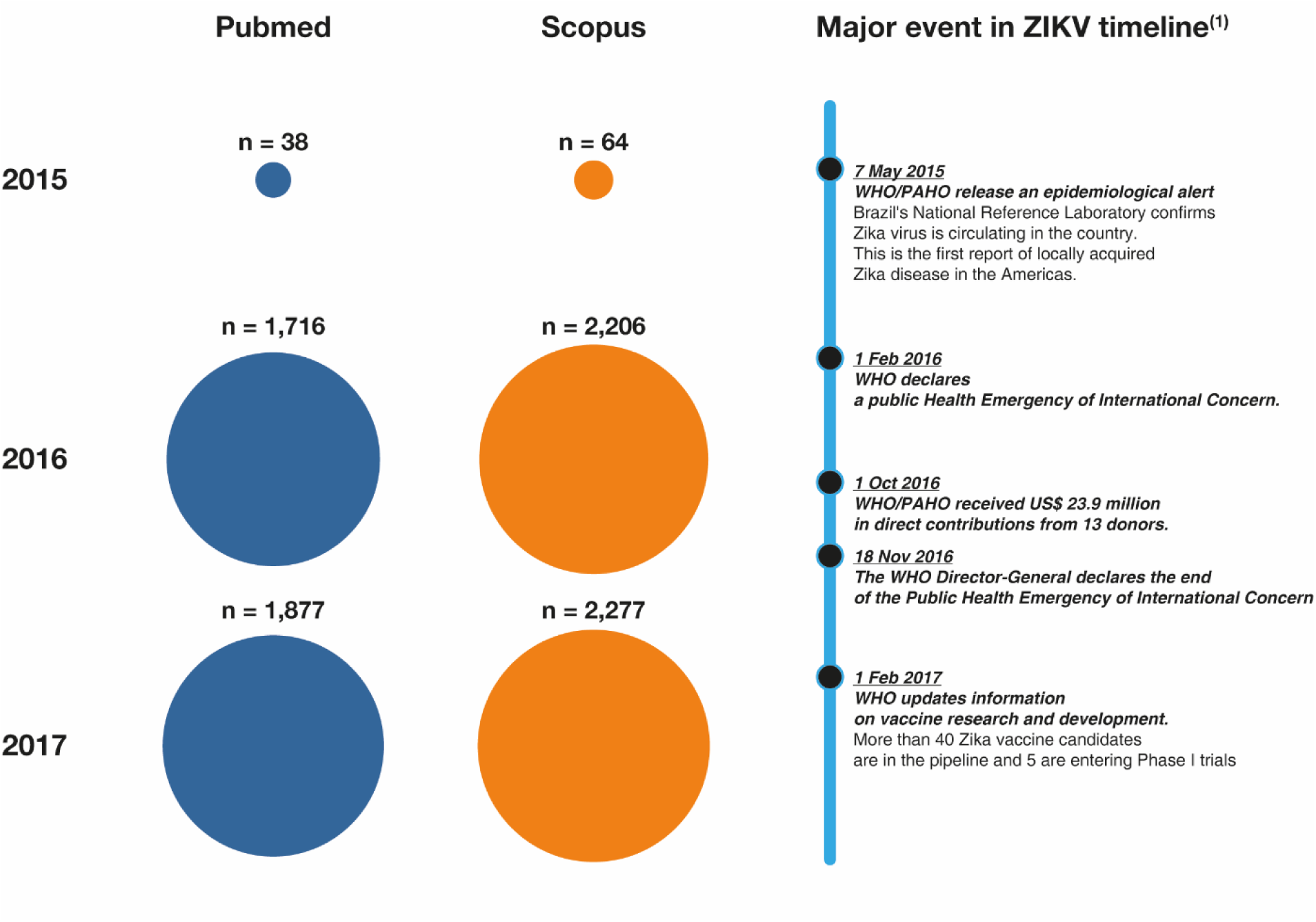
Zika virus associated publications 2015-2017. ^(1)^ The major events in the figure was selected from “WHO - The History of Zika Virus” [23]

This result indirectly showed how active the research has been since the start of the ZIKV pandemic in Americas. However, the time for publication process in journals may vary from months to years. Thus, the publications from 2015 may include the research of yesteryear, but even if we consider the time for the publication process for 2015, there were less than 100 publications. The declaration of a public health emergency by WHO did not only bring public interest to ZIKV research, but also an increase of ZIKV research funding. Accordingly, the number of publications increased dramatically in the past two years.

### ZIKV research field moved from case reports to prospective studies

As a further approach, we decided to generate a bibliometric analysis summary from the Profile RNS website to monitor the trend of ZIKV research. We extracted the data sorted by publication types to investigate the proportion of publication types for the total ZIKV related publications in 2015-2017.

Among various types of publications, we selectively investigated journal articles and case reports. In the case of ZIKV related journal articles, the proportion increased from 35.5% in 2015 (n=27) to 51.3% in 2016 (n=1202), and to 66.8% of journal articles in 2017 (n=1636). On the other hand, the proportion of case reports with ZIKV connections decreased from 14.4% in 2015 (n=11) to 4.3% in 2016 (n=102), and to 1.6% in 2017 (n=40) (Figure 2). In addition, we analyzed the top 20 journals for ZIKV publications during 2015-2017. Most journals where ZIKV related research was published in 2015 had a clinical and surveillance aspect, which shifted to journals that published prospective studies during 2016-2017 (Figure 3).

**Figure 2.**
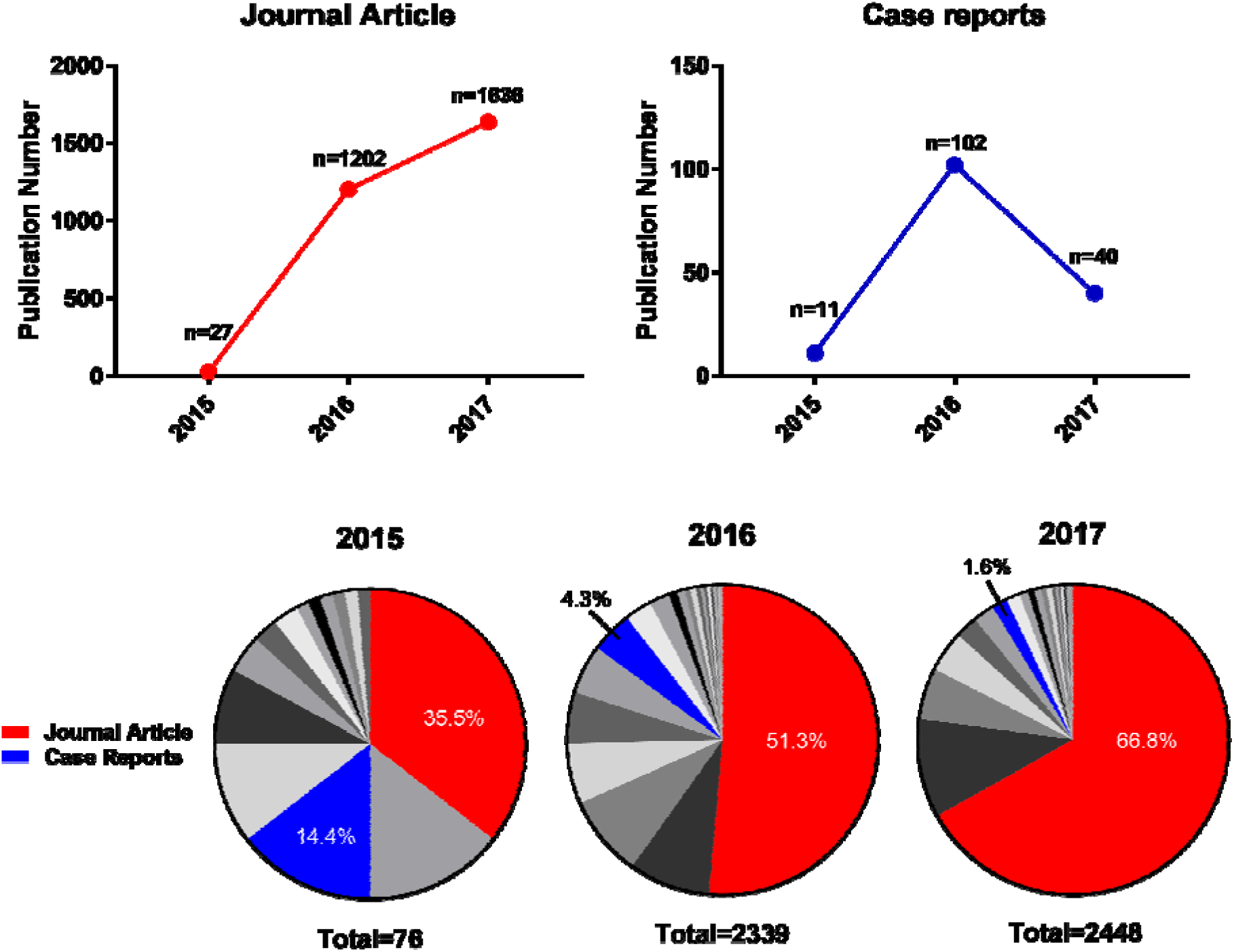
Publication type for Zika virus articles 2015-2017. The publication type was based on the categories assigned to an article in Medline/PubMed. There can be more than one publication type per article, so a single publication might be listed more than once in the table below. Thus, the numbers of publication might add up to more than the total number of publications. The publication types were based on data from Profiles RNS. A full list of types and numbers are available in Additional file 1.

**Figure 3.**
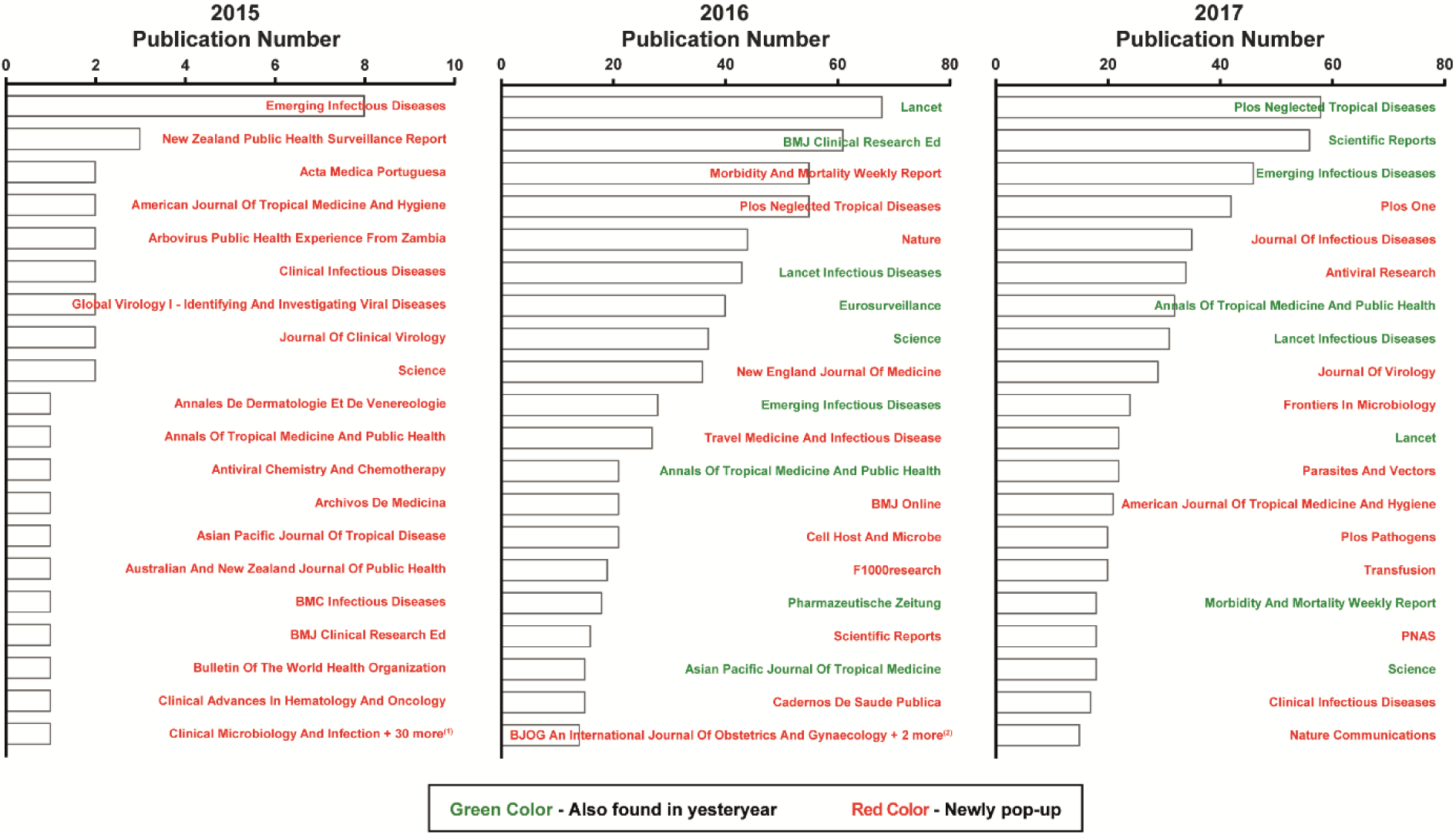
Top 20 journals publishing Zika virus related research 2015-2017. ^(1)^ In 2015, a total of 41 journals had only a single Zika virus (ZIKV) publication. Of the 41 journals with a single ZIKV publication, we selected 11 journals for the figure (by alphabetical order). Since all data in this bibliometric study were collected from 2015, all journals in the 2015 graph were marked as newly pop-up. ^(2)^ In 2016, a total of three journals had fourteen ZIKV publication. Of the three journals with a single ZIKV publication, we selected one journal for the figure (by alphabetical order). A full list of journals is available in Additional file 1.

To find out which articles that were most cited, we extracted the 20 highest cited articles. The result showed that articles about case reports, diagnostic methods, and potential sexual transmission of ZIKV were the most cited in 2015. In 2016, the articles regarding the relationship between ZIKV infection and microcephaly, the relationship between ZIKV infection and Guillain-Barré Syndrome, and development of ZIKV animal model were highly cited. Studies of ZIKV vaccine, ZIKV antiviral treatment, and understanding of ZIKV pathogenicity and tropism were most cited in 2017 (Table 1).

**Table 1.**
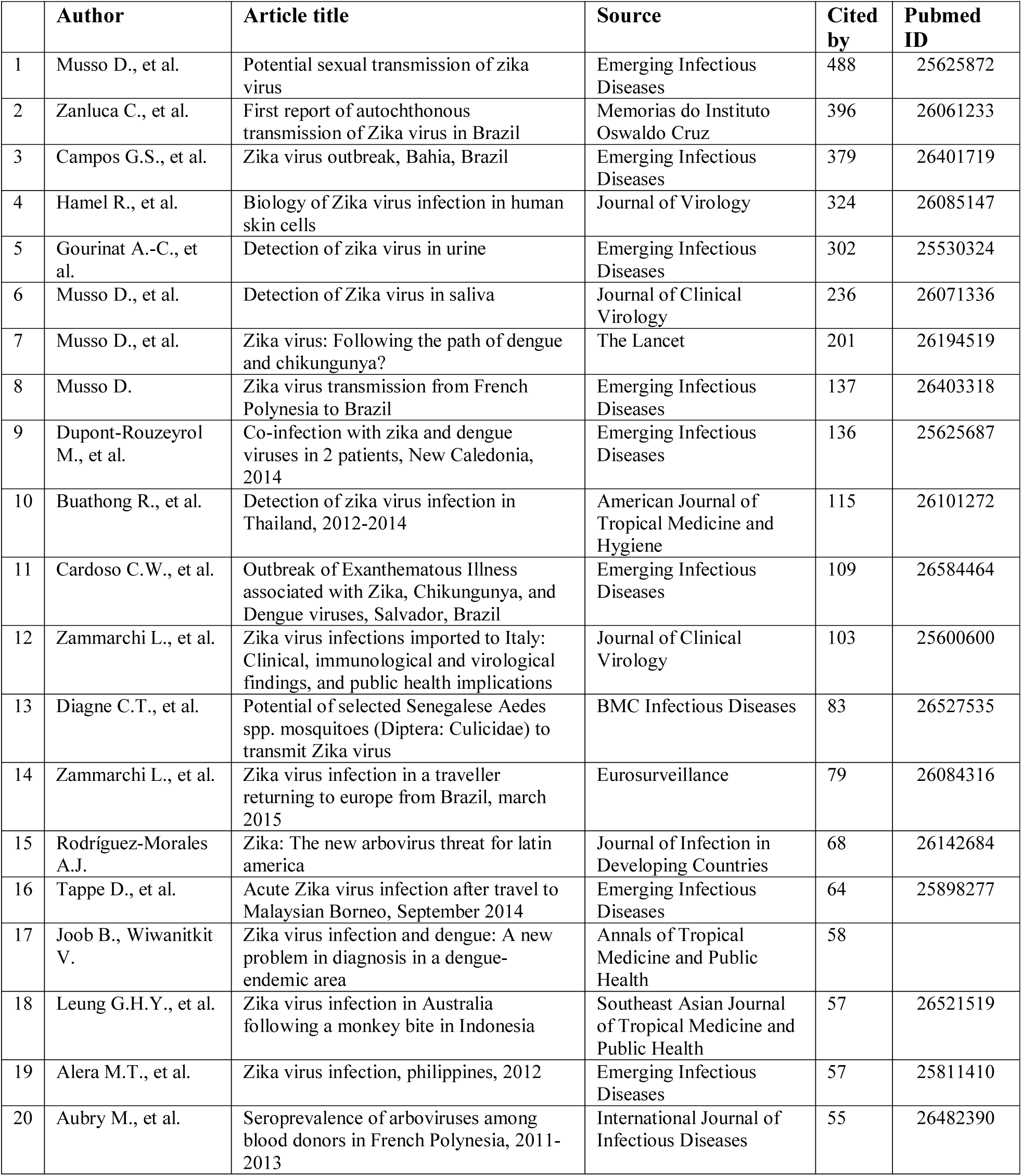
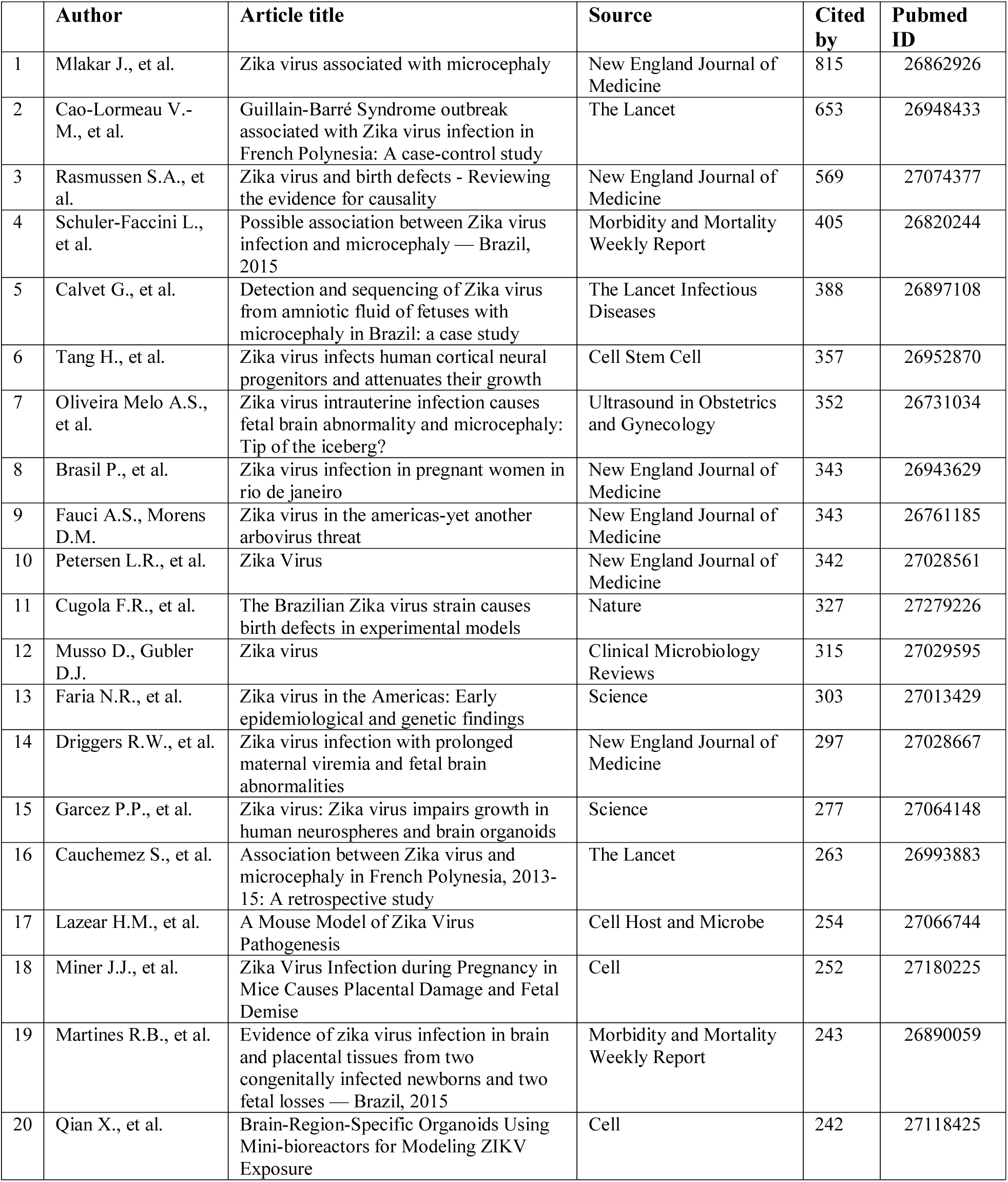
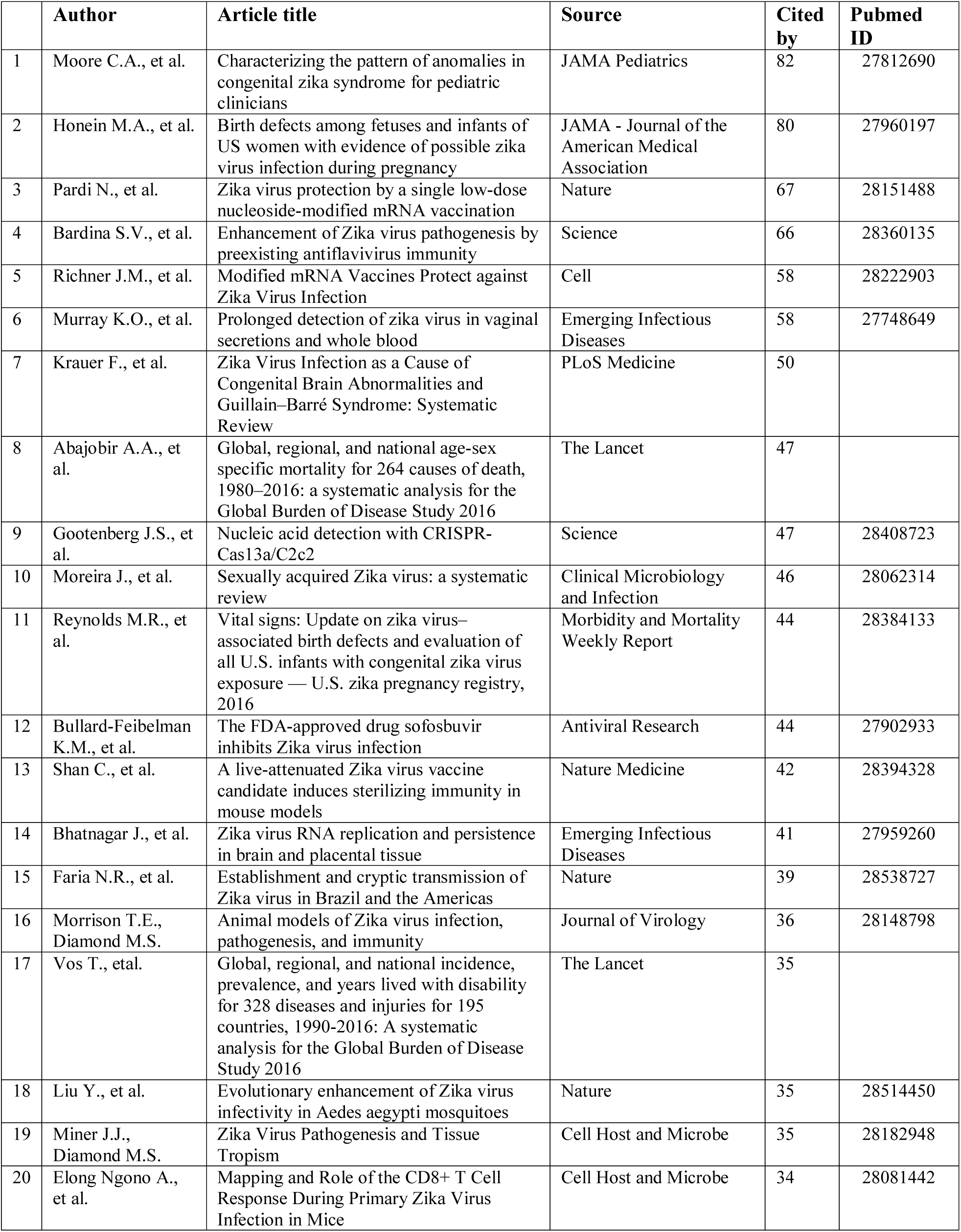
Top 20 cited Zika virus related publications 2015 to 2017.

Overall, when we compared the most cited articles with the pattern of decrease in case reports and increase in total journal articles, we concluded that the ZIKA research trend has shifted from reports of ZIKV case studies and diagnostic methods, to development of ZIKV prevention and treatment, most probably by the gradually accumulated knowledge of ZIKV through the extensive research.

### The geographic distribution of ZIKV publications points to globalization of ZIKV research

We extracted publications by country and generated a world map displaying the geographic distribution of ZIKV publications in 2015-2017. In 2015, 25 countries published ZIKV research, it increased to 111 in 2016 and 139 in the following year, showing that ZIKV research became global during three years (Figure 4).

**Figure 4.**
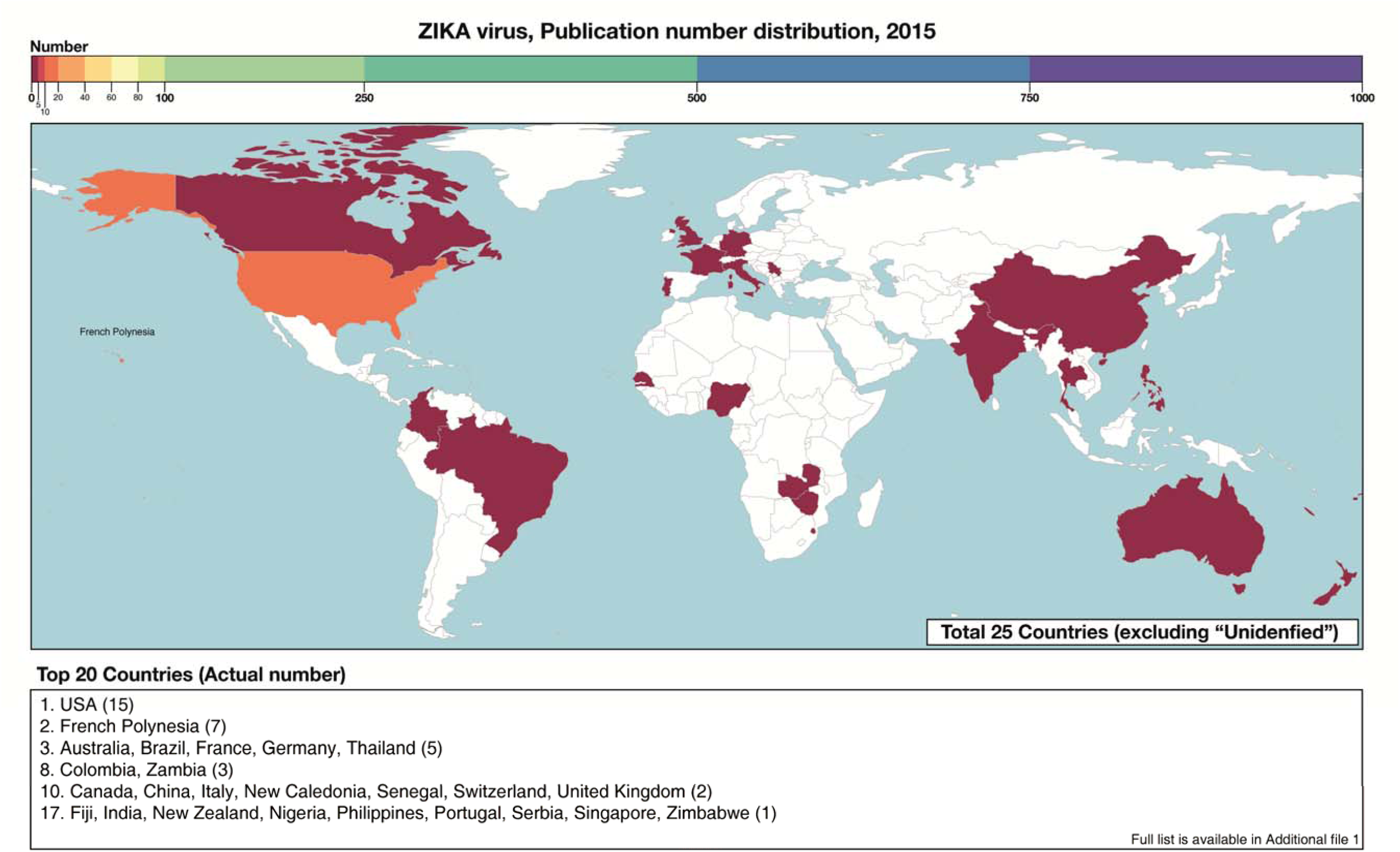
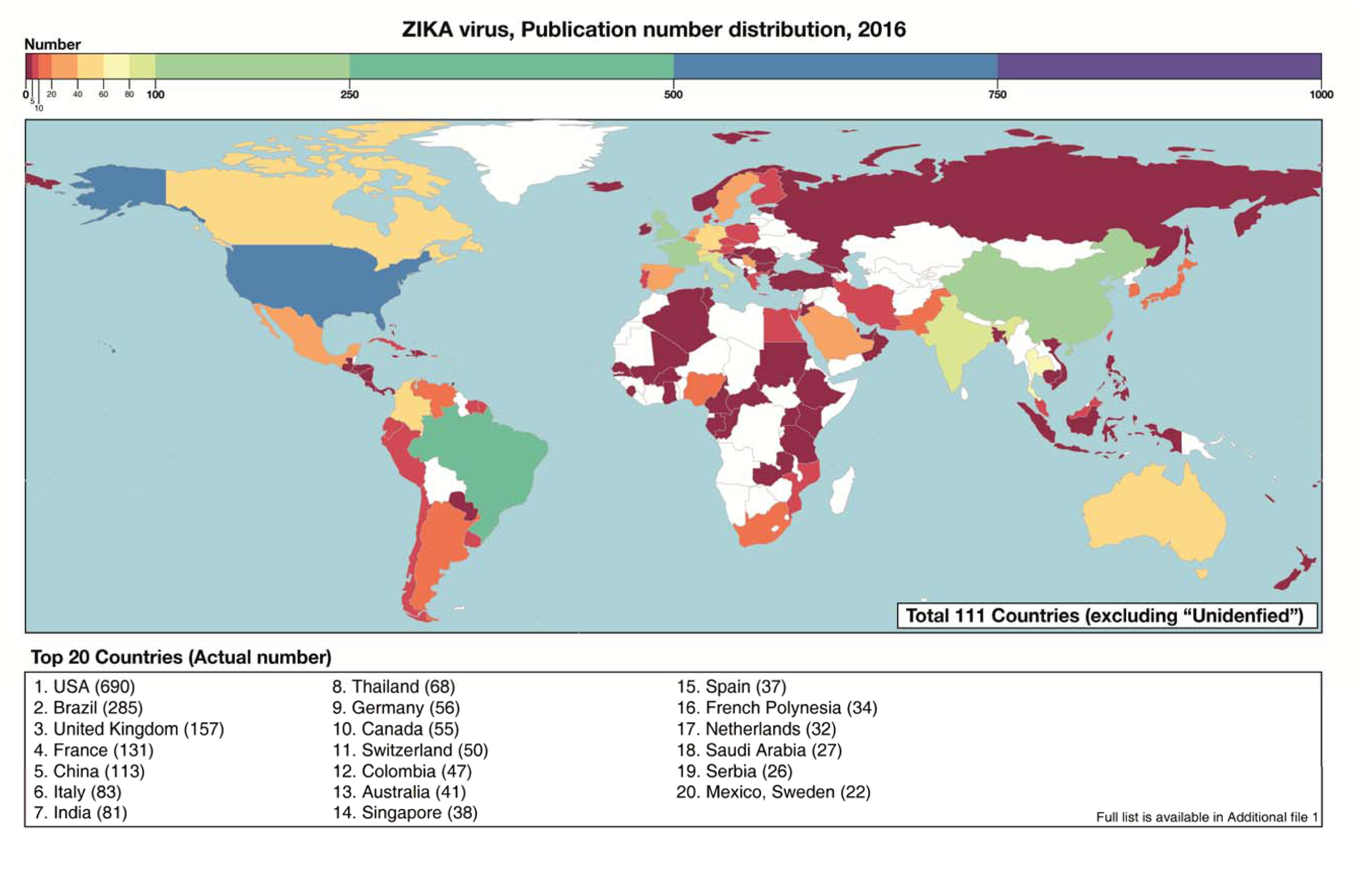
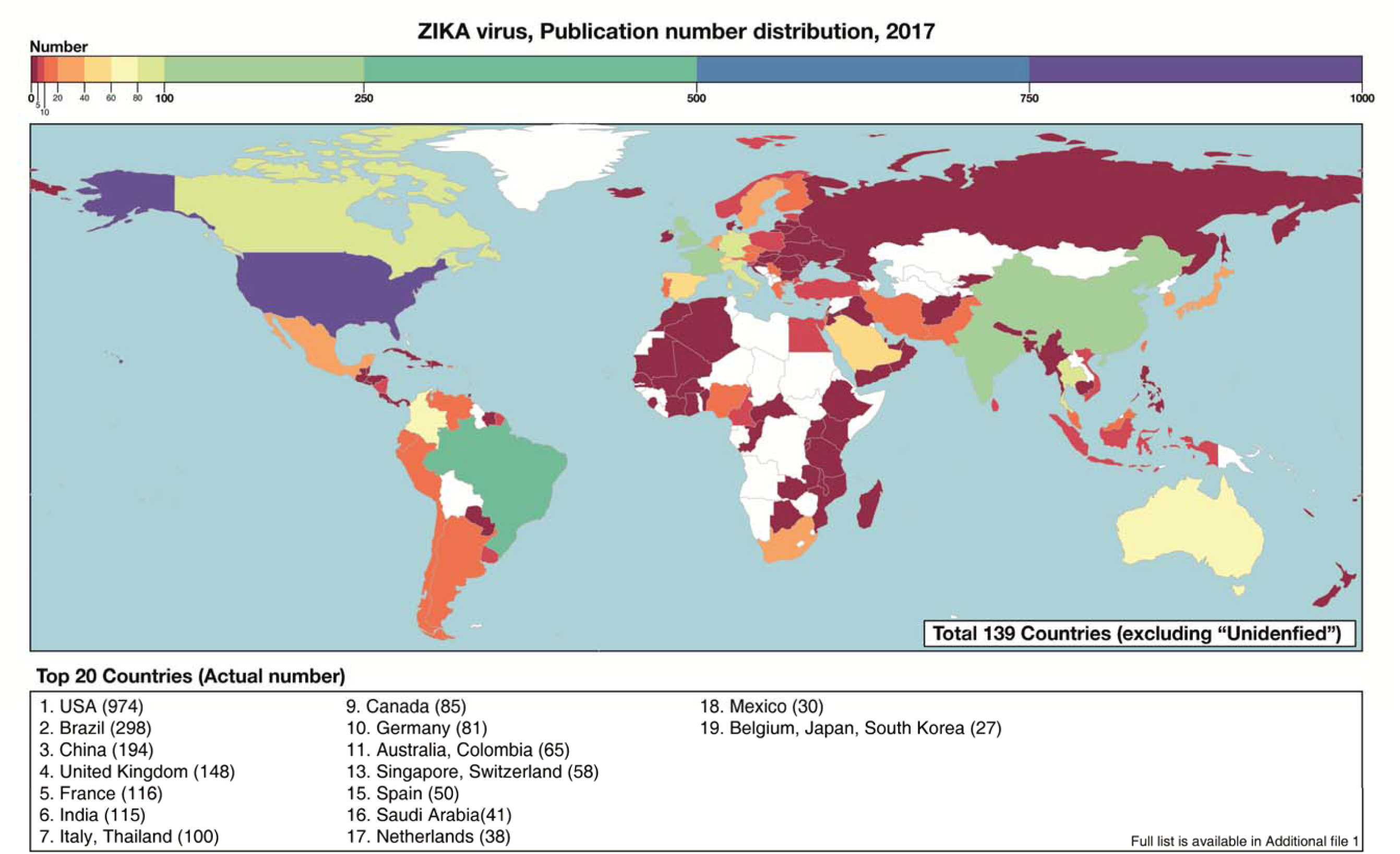
Geographic distribution of Zika virus publications 2015-2017

The United States of America (USA) showed the highest number of ZIKV publications for the whole period 2015-2017, followed by Brazil, the United Kingdom, China, and France. A similar pattern was observed for the top 20 authors with ZIKV related publications, where USA showed the highest number of authors, followed by Brazil, China, and France (Table 2). In addition, a total of 43 authors produced more than 16 articles in 2015-2017 (rank no.1 was 136 articles by Wiwanitkit V).

**Table 2.** Top 20 authors for Zika virus related publications 2015 to 2017.

| Num | Author | Institutes | Publications | Proportion (%) |
| --- | --- | --- | --- | --- |
| 1 | Wiwanitkit, V. | University of Niš, Serbia; Hainan Medical University, China | 136 | 3.00 |
| 2 | Joob, B. | Medical Academic Center, Bangkok, Thailand | 52 | 1.14 |
| 3 | Musso, D. | Institut Louis Malardé, French Polynesia; Aix Marseille Univ, France | 47 | 1.03 |
| 4 | Benelli, G. | University of Pisa, Italy | 46 | 1.01 |
| 5 | Diamond, M.S. | Washington University School of Medicine, USA | 36 | 0.79 |
| 6 | Honein, M.A. | Centers for Disease Control and Prevention, USA | 33 | 0.73 |
| 7 | Shi, P.Y. | University of Texas Medical Branch, USA | 32 | 0.70 |
|  | Weaver, S.C. | University of Texas Medical Branch, USA |  |  |
| 8 | Qin, C.F. | Beijing Institute of Microbiology and Epidemiology, China | 29 | 0.64 |
|  | Vasilakis, N. | University of Texas Medical Branch, USA. |  |  |
| 9 | Govindarajan, M. | Annamalai University, India. | 28 | 0.62 |
|  | Rodriguez-Morales, A.J. | Universidad Tecnológica de Pereira, Colombia. |  |  |
| 10 | Jamieson, D.J. | Emory University, USA | 27 | 0.59 |
| 11 | Meaney-Delman, D. | Centers for Disease Control and Prevention, USA | 26 | 0.57 |
|  | Wiwanitkit, S. | Wiwanitkit House, Thailand |  |  |
| 12 | Gao, G.F. | Chinese Center for Disease Control and Prevention (China CDC), China | 25 | 0.55 |
|  | Leparc-Goffart, I. | French Armed Forces Biomedical Research Institute, France |  |  |
| 13 | Brasil, P. | Instituto Nacional de Infectologia Evandro Chagas, Brazil | 23 | 0.51 |
|  | Cordeiro, M.T. | Aggeu Magalhaes Institute (Instituto Aggeu Magalhães-IAM), Oswaldo Cruz Foundation (Fundação Oswaldo Cruz-FIOCRUZ), Brazil. |  |  |
|  | Fischer, M. | Centers for Disease Control and Prevention, USA |  |  |
|  | Ventura, C.V. | Altino Ventura Foundation, Brazil; HOPE Eye Hospital, Brazil |  |  |
| 14 | McCarthy, M. | Seattle, USA | 22 | 0.48 |
| 15 | Cao-Lormeau, V.M. | Institut Louis Malardé, French Polynesia; Aix Marseille Univ, France | 21 | 0.46 |
|  | Deng, Y.Q. | Beijing Institute of Microbiology and Epidemiology, China |  |  |
|  | Schmidt-Chanasit, J. | Bernhard-Nocht-Institut für Tropenmedizin, Germany; German Centre for Infection Research (DZIF), Germany |  |  |
| 16 | Baud, D. | Department Femme-Mère-Enfant, University Hospital, Switzerland | 20 | 0.44 |
|  | Rasmussen, S.A. | Centers for Disease Control and Prevention, USA |  |  |
| 17 | Oduyebo, T. | Centers for Disease Control and Prevention, USA | 19 | 0.42 |
|  | Rossi, S.L. | University of Texas Medical Branch, USA |  |  |
|  | Shan, C. | University of Texas Medical Branch, USA. |  |  |
|  | Staples, J.E. | Centers for Disease Control and Prevention, USA |  |  |
| 18 | Pierson, T.C. | National Institutes of Health, Bethesda, USA | 18 | 0.40 |
|  | Tanuri, A. | Universidade Federal do Rio de Janeiro, Brazil |  |  |
| 19 | Belfort, R. | Federal University of São Paulo, Brazil | 17 | 0.37 |
|  | Li, X.F. | Beijing Institute of Microbiology and Epidemiology, China |  |  |
|  | Moore, C.A. | University of Colorado, USA |  |  |
|  | Petersen, E.E. | Centers for Disease Control and Prevention, USA |  |  |
|  | Petersen, L.R. | Centers for Disease Control and Prevention, USA |  |  |
|  | Powers, A.M. | Centers for Disease Control and Prevention, USA |  |  |
|  | Rodrigues, L.C. | London School of Hygiene and tropical Medicine, UK |  |  |
| 20 | Grobusch, M.P. | University of Amsterdam, The Netherlands. | 16 | 0.35 |
|  | Hotez, P.J. | National School of Tropical Medicine, Baylor College of Medicine, USA |  |  |
|  | Xie, X. | University of Texas Medical Branch, USA |  |  |

One possible explanation is an increase in ZIKV research funding. In USA, the budget for Ebola virus research was shifted to ZIKV research [24]; the European Union invested funded three research consortia: ZikaPLAN, ZIKAction, ZikAlliance [25]; Brazil, the first site of ZIKV infection in the Americas and host of the World Cup in football in 2016, was provided with emergency research funding [26]. The global support for ZIKV research during this period most probably was a key factor in the increased number of publications globally. It would be interesting to merge the actual budget of ZIKV research in each country for 2015-2017 with the geographic distribution of publications to get a clearer picture for the ZIKV research trend.

## Discussion

Through the bibliometric analysis of Zika related publications, we extracted information describing different factors. The outcome highlighted the importance of gathering public interest to global health issues, and how it can act as a powerful catalyzer to trigger the research field. We also learned that the research efforts resulting from the start of the ZIKV epidemic in the Americas could been seen as an example of global collaboration and efforts to solve a problem when the world is in danger.

After the ZIKV outbreak, the direction of the research started with sharing of information about the risk of the disease, patient cases, and carrying out basic research for the general understanding of the disease. Then it moved to application research based on the understanding from previous basic research, to suggest solutions for the disease (such as vaccines and antiviral treatments). According to the WHO vaccine pipeline tracker, several ZIKV vaccine candidates are currently in clinical phase 1 or 2 [27].

The geographic information showed that during the years after the outbreak of ZIKV in 2015, most countries around the world have taken steps to combat ZIKV and have actively performed research. ZIKV has rapidly become the object of intense investigation by the international research community [20]. However, despite the progress in ZIKV research, many questions remain to be addressed to accelerate the development of effective ZIKV countermeasures

## Conclusions

Through bibliometric analysis of ZIKV related publication for 2015-2017, we have provided an analysis of the ZIKV research trend. Geographic distribution showed that publications from USA, Brazil, and Europe led the scientific production on ZIKV research. It would be interesting to investigate further directions by combining data analyzed by other sociological viewpoints, e.g. the national budget of ZIKV research in different countries. However, we have to remember the importance of support and collaboration from a multidisciplinary effort that is ongoing against the current ZIKV outbreak, to better know how to handle the next inevitable and yet unknown infectious disease threat

